# Neonatal hyperoxia enhances age-dependent expression of SARS-CoV-2 receptors in mice

**DOI:** 10.1101/2020.07.22.215962

**Authors:** Min Yee, E. David Cohen, Jeannie Haak, Andrew M. Dylag, Michael A. O’Reilly

**Affiliations:** The Department of Pediatrics, School of Medicine and Dentistry, The University of Rochester, Rochester, NY 14642

**Author notes:** Address Correspondence to: Michael A. O’Reilly, Ph.D., Department of Pediatrics, Box 850, The University of Rochester, School of Medicine and Dentistry, 601 Elmwood Avenue, Rochester NY 14642.

**Keywords:** Angiotensin Converting Enzyme 2, COVID-19, Hyperoxia, Mice, Transmembrane protease/serine subfamily member 2

## Abstract

The severity of COVID-19 lung disease is higher in the elderly and people with pre-existing co-morbidities. People who were born preterm may be at greater risk for COVID-19 because their early exposure to oxygen at birth increases their risk of being hospitalized when infected with RSV and other respiratory viruses. Our prior studies in mice showed how high levels of oxygen (hyperoxia) between postnatal days 0-4 increases the severity of influenza A virus infections by reducing the number of alveolar epithelial type 2 (AT2) cells. Because AT2 cells express the SARS-CoV-2 receptors angiotensin converting enzyme (ACE2) and transmembrane protease/serine subfamily member 2 (TMPRSS2), we expected their expression would decline as AT2 cells were depleted by hyperoxia. Instead, we made the surprising discovery that expression of *Ace2* and *Tmprss2* mRNA increases as mice age and is accelerated by exposing mice to neonatal hyperoxia. ACE2 is primarily expressed at birth by airway Club cells and becomes detectable in AT2 cells by one year of life. Neonatal hyperoxia increases ACE2 expression in Club cells and makes it detectable in 2-month-old AT2 cells. This early and increased expression of SARS-CoV-2 receptors was not seen in adult mice who had been administered the mitochondrial superoxide scavenger mitoTEMPO during hyperoxia. Our finding that early life insults such as hyperoxia enhances the age-dependent expression of SARS-CoV-2 receptors in the respiratory epithelium helps explain why COVID-19 lung disease is greater in the elderly and people with pre-existing co-morbidities.

## INTRODUCTION

COVID-19 is an infectious disease of the lung caused by the severe acute respiratory syndrome coronavirus (SARS-CoV-2). As of July 2020, the World Health Organization reported this virus has infected more than 10 million people worldwide and killed approximately 500,000 people (https://covid19.who.int). Common symptoms include fever, cough, fatigue, shortness of breath, and loss of olfactory or gustatory function. While the majority of cases are mild, some people progress into severe acute respiratory distress syndrome, multi-organ failure, thrombosis, and septic shock. The severity of disease and mortality is highest among the elderly and people who have pre-existing lung or heart disease. There is growing evidence that asymptomatic children and young adults with COVID-19 may be at risk for heart disease, inflammatory vascular disease, and stroke ^1^. People who were born preterm may be at great risk for COVID-19 because they are already at risk for hospitalization following infection with RSV, rhinovirus, human bocavirus, metapneumovirus, and parainfluenza viruses ^2^. They may also develop pulmonary vascular disease and heart failure ^3,4^, autism-like disorders ^5,6^, and retinopathy ^7^ that puts them at further risk for COVID-19. Identifying mechanisms that drive susceptibility to pandemic viral infections like SARS-CoV-2 is therefore of great concern to susceptible individuals and their families.

The severity of COVID-19 is likely to be related to age-related changes in SARS-CoV-2 receptors and how the immune system responds to infection ^1^. Emerging evidence indicates high-risk individuals with SARS-CoV-2 have high rates of alveolar epithelial type 2 (AT2) cell infection, suggesting disease severity may be related to higher alveolar expression of the SARS-CoV-2 receptor angiotensin converting enzyme (ACE2) and its co-receptor transmembrane protease/serine subfamily member 2 (TMPRSS2) ^8,9^. In fact, a recent meta-analysis of 700 people with predicted COVID-19 co-morbidities found that their lungs expressed high levels *Ace2* mRNA ^10^. ACE2 is a zinc containing metalloprotease present at the surface of cells in the lung, heart, intestines, kidneys, and brain. It lowers blood pressure by catalyzing the hydrolysis of the vasoconstrictive molecule angiotensin II to angiotensin (1-7). ACE2 co-precipitates with transmembrane protease/serine subfamily member 2 (TMPRSS2) which hydrolyzes the S protein on coronaviruses, thus enabling viral entry into infected cells ^9,11^. Higher expression of these proteins in AT2 cells would theoretically lead to higher rates of infection in the distal lung. Infected AT2 cells produce inflammatory mediators that could contribute to a lethal cytokine storm ^12,13^. They may also die. Loss of AT2 cells below a critical threshold could compromise alveolar homeostasis because they produce surfactant and serve as adult stem cells for the alveolar epithelium ^14^. In fact, high rates of AT2 infection have been seen in people who have succumbed to H5N1, a highly pathogenic avian strain of influenza A virus ^15-17^. But whether aging or pre-existing lung co-morbidities like preterm birth enhance the severity of respiratory viral infections via changing expression of viral receptors is not yet known.

Since preterm infants are exposed too soon to oxygen, we have been using mice to understand how high levels of oxygen at birth increases the severity of influenza A virus infection in adults. We previously reported how adult mice exposed to hyperoxia (100% oxygen) between postnatal days 0-4 develop persistent inflammation and fibrotic lung disease when infected with influenza A viruses HKx31 (H3N2) or PR8 (H1N1) ^18,19^. Neonatal hyperoxia does not enhance primary infection ^20^ or clearance ^21^ of the virus. Instead, it reduced the number of adult AT2 cells by ∼50%, thus lowering the number available to maintain alveolar homeostasis and epithelial regeneration after infection ^22^. Because neonatal hyperoxia reduces the number of AT2 cells, we predicted it would reduce the alveolar expression of ACE2 and TMPRSS2 in the lung. Instead, we made the surprising discovery that expression of ACE2 and TMPRSS2 increases as mice age and this age-dependent expression can be enhanced by early exposure to hyperoxia. Our findings in mice suggest temporal and spatial changes in expression of SARS-CoV-2 receptors may contribute to the increased severity of COVID-19 seen in the elderly and people with pre-existing co-morbidities, including those born preterm.

## RESULTS

### ACE2 is initially expressed by Club cells and then by AT2 cells as mice age

The localization of ACE2 was examined in the lungs of mice between PND4 and 2 years of age by immunohistochemistry so as to better understand the temporal spatial pattern of its expression. ACE2 was primarily detected in airway epithelial cells with minimal staining seen in the alveolar space (**Figure 1a**). The intensity of ACE2 staining increased steadily in the airway epithelium throughout the life of the mouse. A rare ACE2-positive alveolar cells (arrows) was first observed on PND7 and then steadily increased in number between 6 and 24 months of age. Western blotting for ACE2 confirmed that the abundance of ACE2 protein became progressively enriched in the whole lungs of 12- and 24-month-old mice relative to those of mice harvested at 2 months of age (**Figure 1b**). ACE2 mRNA levels were similarly increased in the whole lungs of 24-month-old mice than in those of mice harvested at 2 months of age (**Figure 1c**).

**Figure 1.**
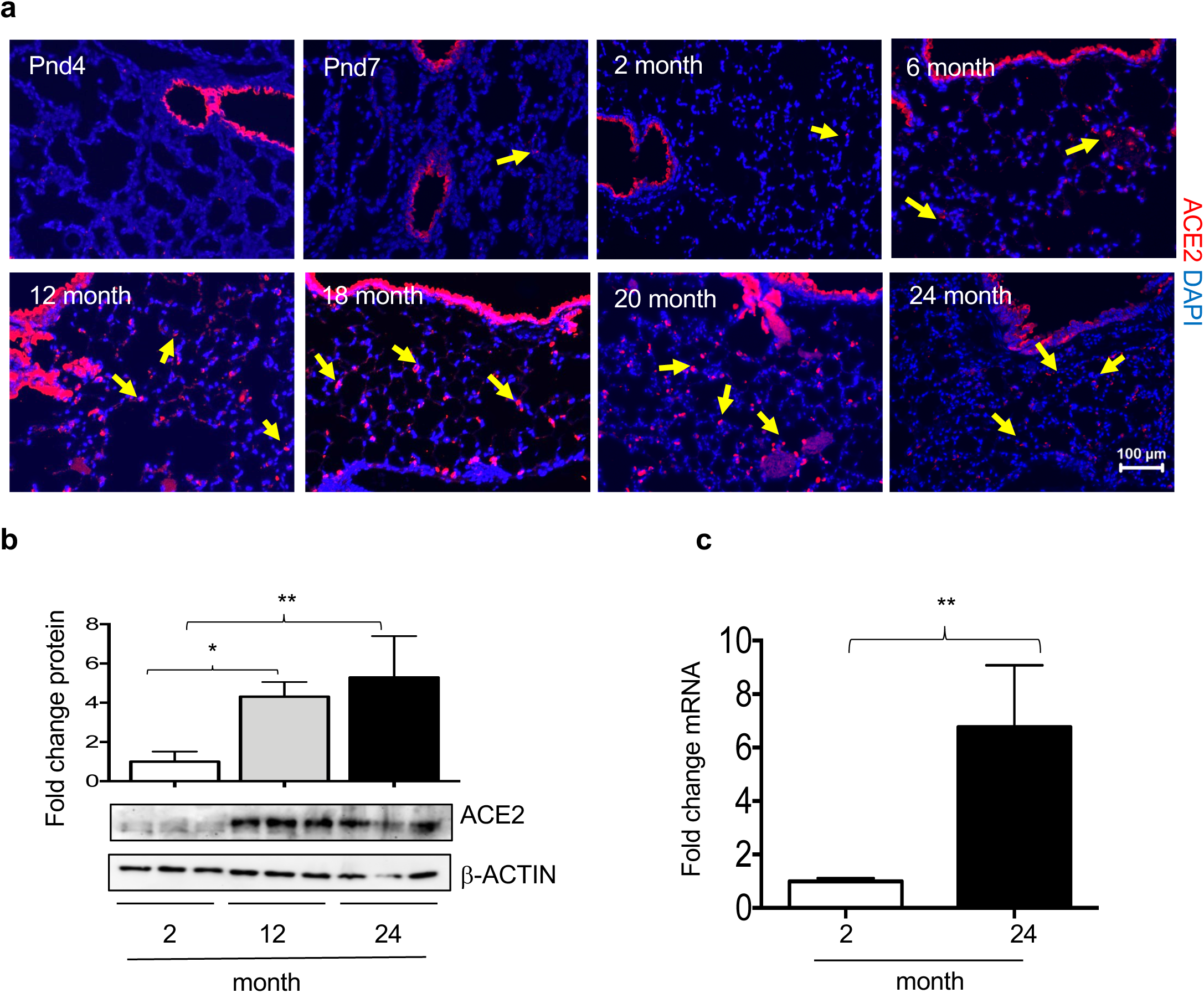
ACE2 expression changes in lung as mice age. (**a**) Lungs harvested from mice of different ages were stained for ACE2 (red) and counterstained with DAPI (blue). ACE2 was detected in airways of all mice and alveolar regions (yellow arrows). Bar = 100 μm. (**b**) Lungs homogenates prepared from 2-month, 12-month, and 24-month-old mice were immunoblotted for ACE2 and β-ACTIN as a loading control. Each lane represents an individual mouse. Band intensity of ACE2 to β-ACTIN was quantified and graphed as fold change relative to 2-month samples. Bars reflect mean ± SD graphed. (**c**) qRT-PCR was used to quantify *Ace2* mRNA in total lung homogenates of 2-month and 24-month-old mice. Data is graphed as the fold change of *Ace2* after normalizing to *18S* RNA. Bars reflect mean ± SD graphed as fold change over 2-month values. Statistical significance is comparisons for all pairs using Tukey-Kramer HSD test, with *P≤0.05; **P≤0.01.

Co-staining with antibodies for ACE2 and the Club cell marker secreteglobin1a1 (Scgb1a1) showed extensive co-localization along the airways at both 2 and 12 months of age (**Figure 2a**), but the intensity of ACE2 staining was significantly higher at 12 months of age than at 2 months of age (**Figure 2b**). Co-staining for ACE2 and the AT2 cell marker proSP-C revealed that the vast majority of ACE2+ cells in the alveoli were AT2 cells (**Figure 2c**). Approximately 20% of proSP-C+ AT2 cells expressed ACE2 at 2 months while 80% of proSP-C+ AT2 cells expressed it at 12 months (**Figure 2d**). These findings reveal that ACE2 is primarily expressed by the airway Club cells of young adult mice but becomes increasingly expressed by AT2 cells as mice age.

**Figure 2.**
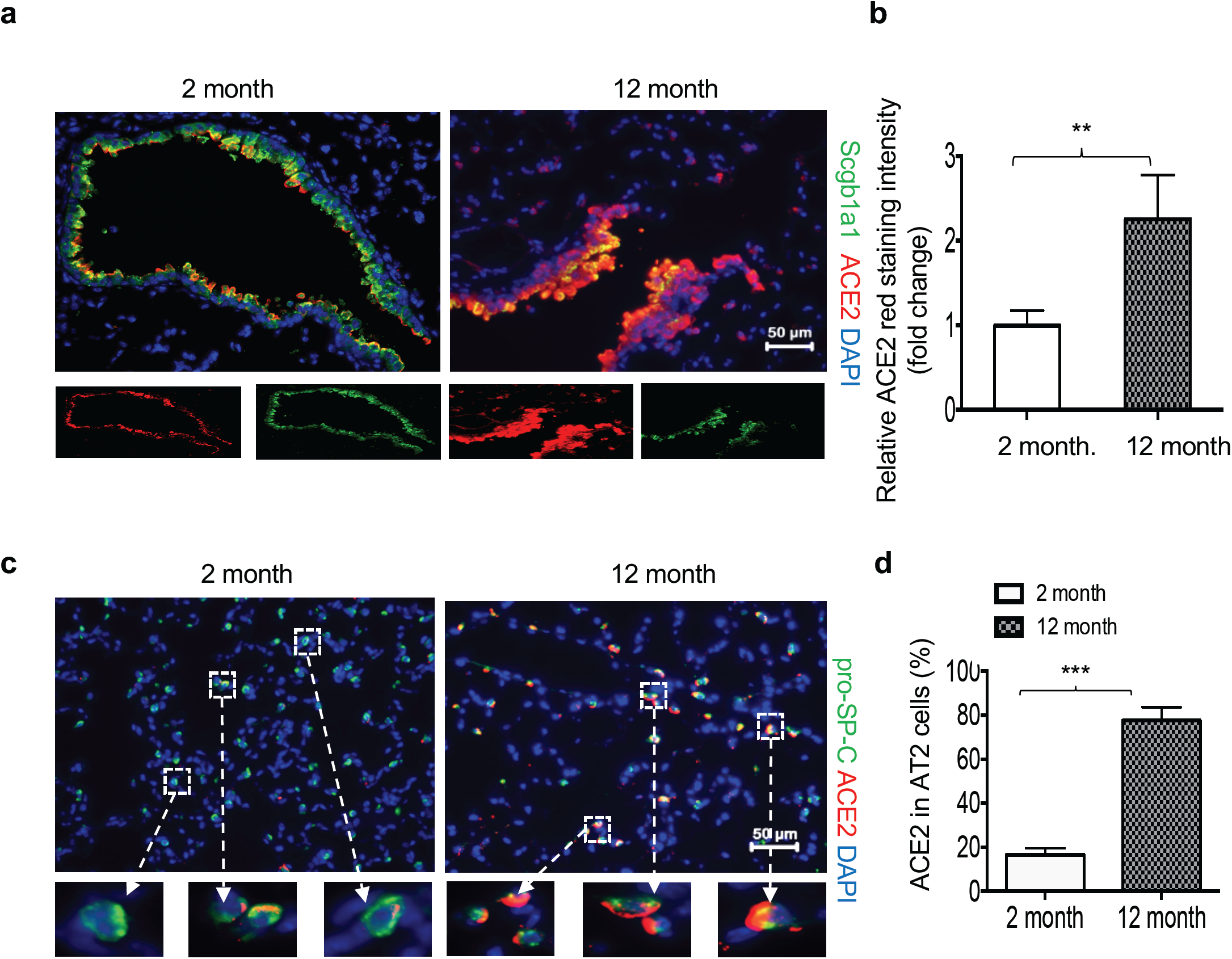
Aging increases ACE2 expression in airway Club and alveolar type 2 cells. (**a**) Lungs from 2-month and 12-month-old mice were immunostained for ACE2 (red), Scgb1a1 (green), and counterstained with DAPI (blue). Boxed sections are individual ACE2 and Scgb1a1 stains. (**b**) Quantitation of ACE2 Red staining intensity. All the cells were imaged using identical exposure time. Scale bar = 50 μm. (**c**) Lungs were stained for ACE2 (red), proSP-C (green), and counterstained with DAPI (blue). Boxed sections are enlarged below each figure. (**d**) The proportion of proSP-C+ cells expressing ACE2 was quantified and graphed. Statistical significance is comparisons for all pairs using Tukey-Kramer HSD test, with **P≤0.01; ***P≤0.001. Bar = 50 μm.

### Neonatal hyperoxia enhances the age-dependent changes in ACE2 expression

We previously showed that adult mice exposed to 100% oxygen between PND0-4 (**Figure 3a**) have fewer AT2 cells than mice exposed to room air ^23^and thus expected ACE2 expression to be lower in the lungs of mice exposed to neonatal hyperoxia than in those of controls. It was therefore surprising to find that the levels of ACE2 protein were higher in the lungs of 2-month-old mice that were exposed to neonatal hyperoxia than in age-matched control lungs (**Figure 3b**). The levels of Ace2 mRNA were also increased in the lungs of neonatal hyperoxia-exposed mice at 2 months of age and remained higher than in the lungs of age-matched controls at 6 and 12 months of age (**Figure 3c**). To determine the amount of oxygen needed to stimulate the expression of *Ace2*, the lungs of 2-month-old mice exposed to 0, 40, 60 or 80% oxygen from PND0-4 were examined by qRT-PCR (**Figure 3d**). While 40% oxygen was not sufficient to induce *Ace2* mRNA, the levels of *Ace2* expression was significantly higher in mice exposed to 60% and 80% oxygen relative to controls. Exposing mice to a low chronic dose of oxygen (40% for 8 days) that does not alter alveolar development ^24^ also failed to increase Ace2 levels relative to controls (data not shown). Because 40% oxygen for 8 days is higher cumulative dose of oxygen than 60% for 4 days, these findings suggest that oxygen alone may not be stimulating *Ace2* expression.

**Figure 3.**
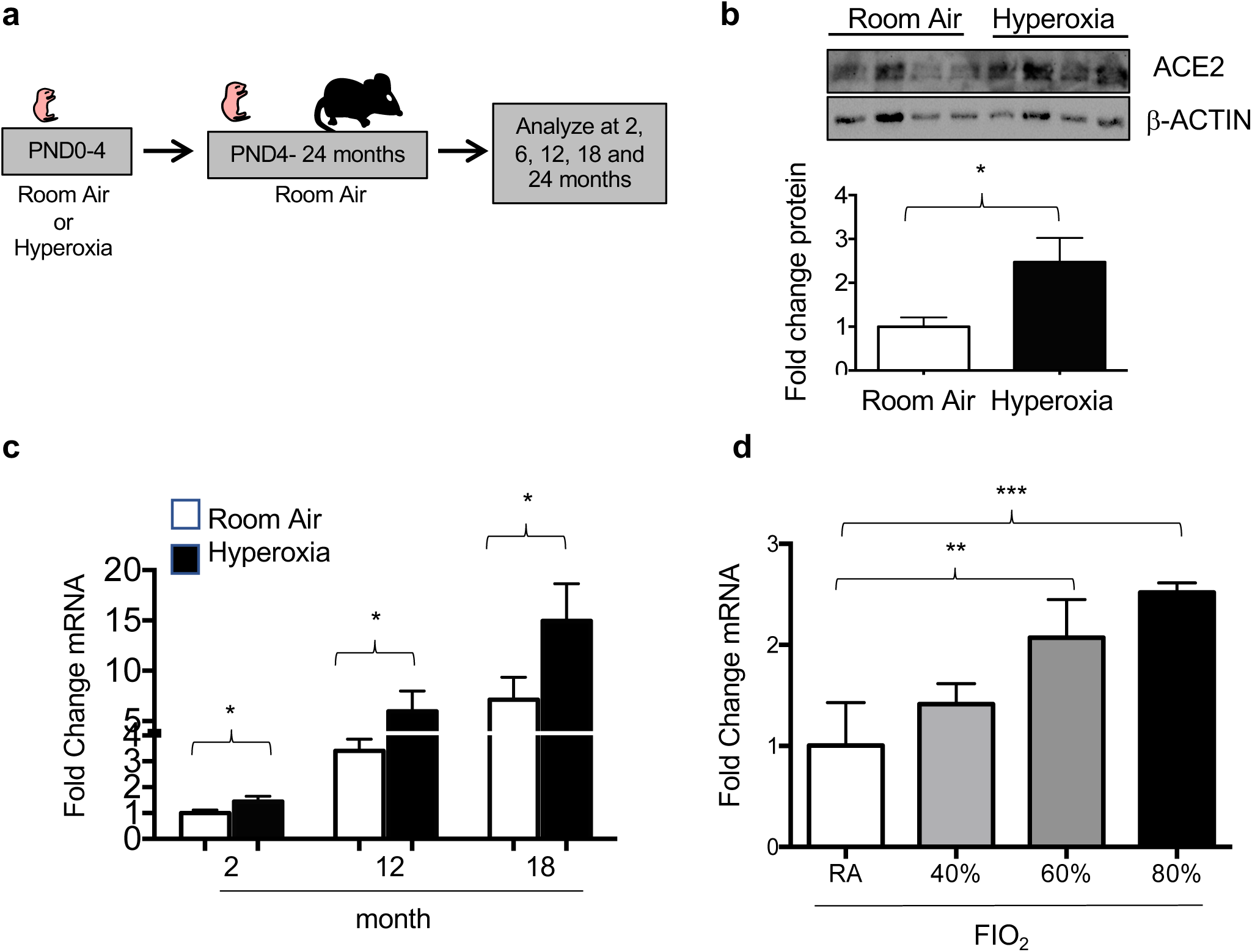
Neonatal hyperoxia stimulates expression of ACE2 in adult mice. (**a**) Cartoon showing the experimental approach of exposing newborn mice to hyperoxia. (**b**) Total lung homogenates were immunoblotted for ACE2 and β-ACTIN as a loading control. Data is graphed as mean ± SD fold change over room air values. (**c**) qRT-PCR was used to quantify *Ace2* mRNA in total lung homogenates of 2-, 12-, and 18-month-old mice exposed to room air or hyperoxia between PND0-4. Values were normalized to expression of *18S* RNA and graphed as mean ± SD fold change of ACE2 in 2-month-old room air mice. (**d**) qRT-PCR was used to quantify *Ace2* mRNA in total lung homogenates of 2-month-old mice exposed to room air, 40%, 60%, or 80% oxygen between PND0-4. Values were normalized to expression of *18S* RNA and graphed as fold change of ACE2 in 2 month room air mice. Statistical significance is comparisons for all pairs using Tukey-Kramer HSD test with *P≤0.05; **P≤0.01; ***P≤0.001.

Immunohistochemistry was used to further understand how hyperoxia affected ACE2 expression in the adult lung. While neonatal hyperoxia increased intensity of ACE2 staining in the airway, it most obviously increased the number of alveolar cells with detectable ACE2 (**Figure 4a**). When quantified, neonatal hyperoxia increased the number of alveolar cells expressing ACE2 by approximately 50% at 2, 6 and 12 months of age (**Figure 4b**). The increased alveolar expression seen at 2 months of age was primarily attributed to increased expression by proSP-C+ AT2 cells; however, this difference resolved at 6 and 12 months of age as more AT2 cells in control lungs began to express ACE2 (**Figure 4c**).

**Figure 4.**
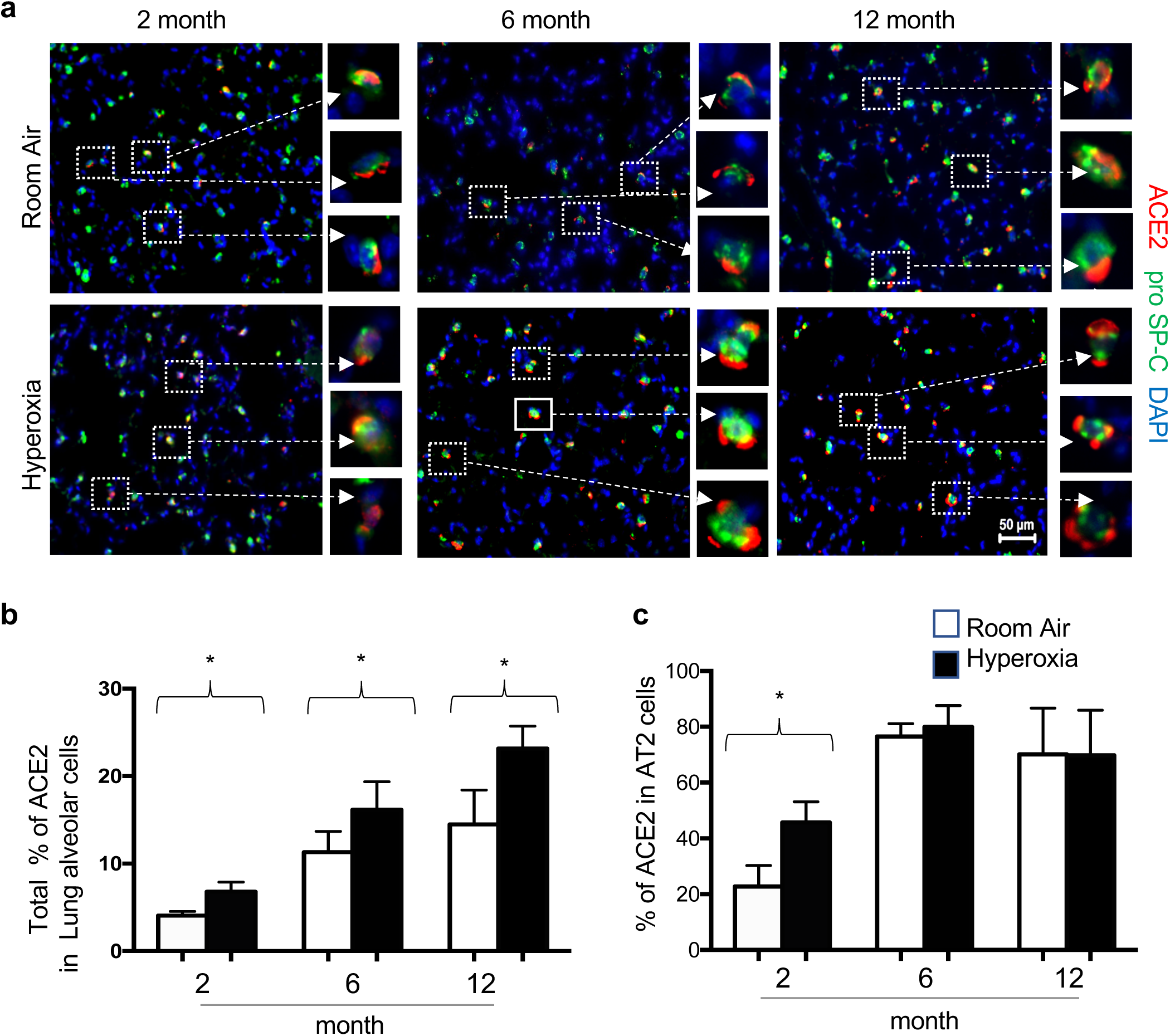
Neonatal hyperoxia stimulates expression of ACE2 in alveolar type 2 cells. (**a**) Lungs of 2-, 6- and 12-month-old mice exposed to room air or hyperoxia between pnd0-4 were stained for ACE2 (red), proSP-C (green), and DAPI. Upper rows reflect room air and lower rows reflect hyperoxia between PND0-4. Boxed regions are enlarged to the right of each image. (**b**) The proportion of ACE2-positive to total DAPI cells was quantified and graphed. (**C**) The proportion of proSP-C+ cells that express ACE2 were quantified and graphed. Values in b, c represent mean ± SD of 4-5 lungs per group with stated P values in the graphs. Statistical significance is comparisons for all pairs using Tukey-Kramer HSD test with*P≤0.05.

### Anti-oxidants block oxygen-dependent changes in ACE2 expression

Prior studies by us and other investigators showed that administering the mitochondrial superoxide scavenger mitoTEMPO to mice during exposure to hyperoxia (**Figure 5a**) prevents the alveolar simplification and cardiovascular disease observed when these mice reach adulthood ^25-27^. qRT-PCR revealed administering mitoTEMPO during hyperoxia blunted the oxygen-dependent increase in *Ace2* mRNA seen in 2-month-old mice (**Figure 5a, b**). Immunohistochemistry confirmed mitoTEMPO reduced the number of AT2 cells with detectable levels of ACE2 protein (**Figure 5c, d**). It also reduced the intensity of ACE2 staining in airway Club cells (**Figure 5e, f**). Interestingly, while mitoTEMPO did not affect ACE2 staining in control mice, it reduced the numbers of alveolar ACE2+ cells in the lungs of hyperoxia-exposed mice lower than controls.

**Figure 5.**
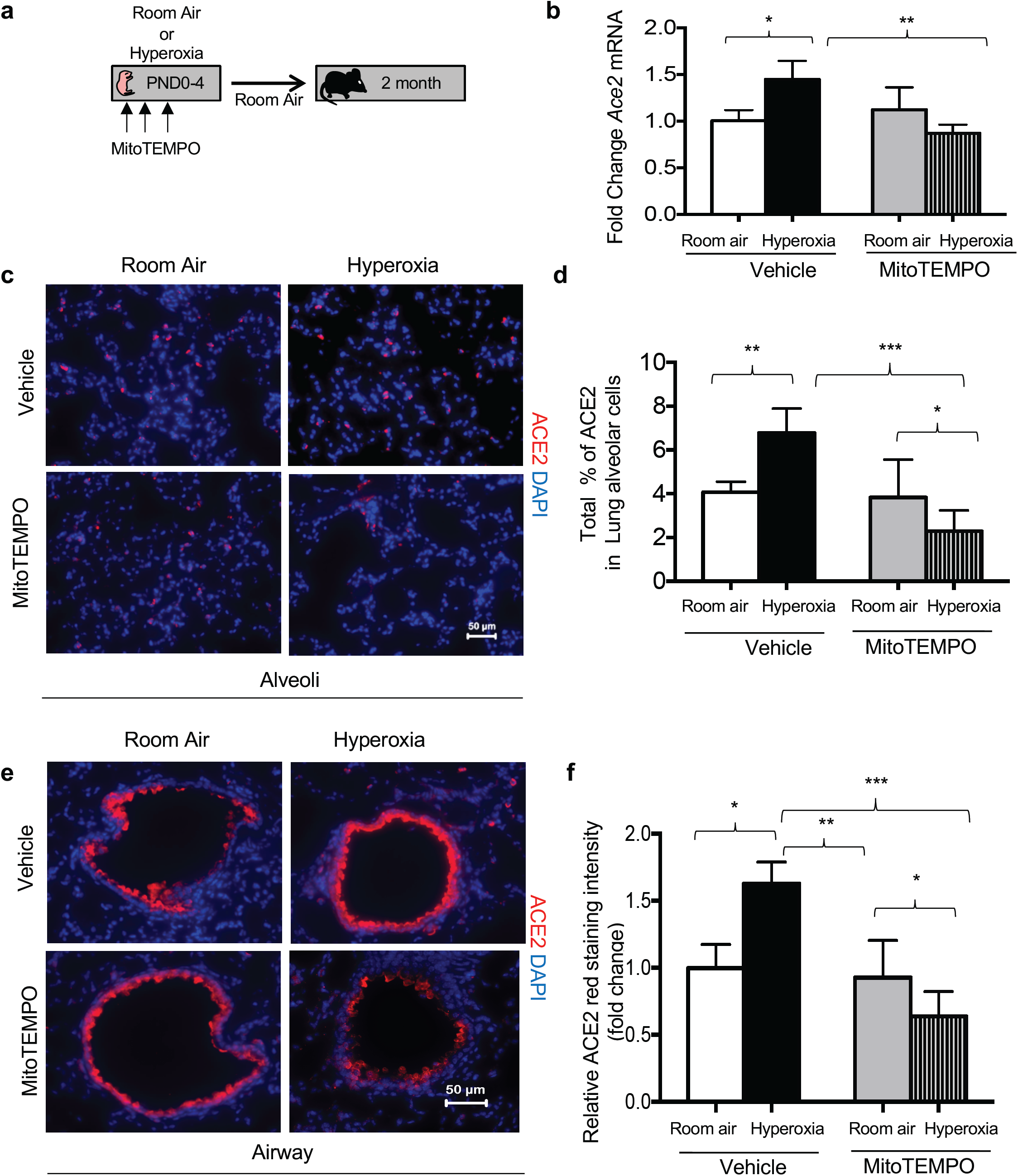
Anti-oxidants prevent hyperoxia from stimulating expression of ACE2. (**a**) Cartoon showing the experimental approach of exposing newborn mice to hyperoxia and treated with mitoTEMPO (d1-d3). (**b**) qRT-PCR was used to measure *Ace2* mRNA expression in 2-month-old mice exposed to room air or hyperoxia as vehicle or mitoTEMPO between PND0-4. Values reflect mean ± SD of 4-5 mice per group and graphed as fold change over mice administered room air and vehicle control. Expression of *Ace2* mRNA was normalized to *18S* rRNA and mean ± SD values graphed relative to room air values. (**c**) Lung alveoli were stained for ACE2 (red), and counterstained with DAPI (blue). (**d**) Total % of ACE2 cells in lung alveoli. (**e**) Lung airways were stained for ACE2 (red), and counterstained with DAPI (blue). (**f**) Quantitation of ACE2 Red staining intensity. All the cells were imaged using identical exposure time. Scale bar = 50 μm; Quantitation of ACE2 Red was derived from images. Statistical significance is comparisons for all pairs using Tukey-Kramer HSD test with *P≤0.05; **P≤0.01; ***P≤0.001.

### Neonatal hyperoxia stimulates age-dependent changes in TMPRSS2

TMPRSS2 is an endoprotease expressed by respiratory epithelial cells that facilitates viral entry of coronaviruses into epithelial cells ^9^. The levels of Tmprss2 mRNA and protein were examined in the lungs of 2-, 12- and 18-month-old mice that were exposed to neonatal hyperoxia and room air from PND0-4 by qRT-PCR and western blotting. *Tmprss2* mRNA was readily detected in the lungs of 2-month-old mice, and increased ∼5-fold at 12 months and ∼8-fold at 18 months (**Figure 6a**). Neonatal hyperoxia further increased *Tmprss2* expression by ∼50% at each time-point examined. Western blotting similarly showed that the levels of TMPRSS2 protein were higher in the whole lung lysates of mice exposed to neonatal hyperoxia than in those of control mice (**Figure 6b**). As observed for *Ace2* expression, exposure to ≥ 60% oxygen from PND4-0 was required to significantly increase the levels of Tmprss2 mRNA in the lungs of mice at 2 months of age (**Figure 6c**). Exposure to 40% oxygen from PND0-8 also failed to change Tmprss2 expression in adult mice (data not shown) while the administration of mitoTEMPO to mice during exposure blunted the effects of neonatal hyperoxia on Tmprss2 mRNA (**Figure 6d**). Together, these findings suggest age and neonatal hyperoxia have similar effects on increasing TMPRSS2 as they do for ACE2.

**Figure 6.**
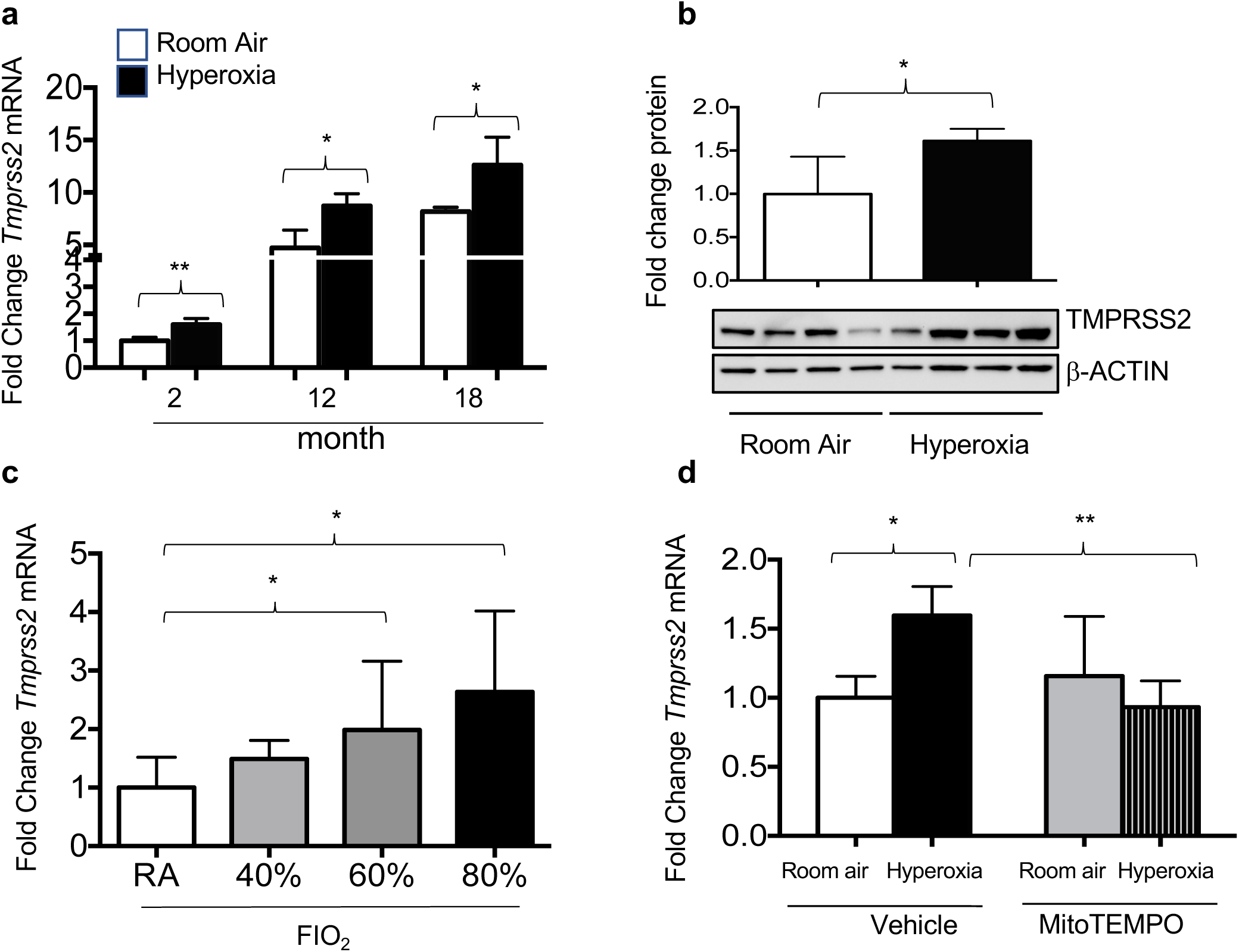
Neonatal hyperoxia stimulates age-dependent expression of *Tmprss2* mRNA. (**a**) qRT-PCR was used to quantify *Tmprss2* mRNA in total lung homogenates of 2-, 12-, and 18-month-old mice exposed to room air or hyperoxia between PND0-4. Values were normalized to expression of *18S* RNA and graphed as fold change of ACE2 in 2-month-old room air mice. (**b**) Western blot-based quantification of TMPRSS2. Data in panels A-D reflect mean ± SD and graphed as fold change relative to control mice exposed to room air. (**c**) qRT-PCR was used to measure *Tmprss2* mRNA in total lung homogenates of 2 month mice exposed to room air, 40%, 60%, or 80% oxygen between PND0-4. (**d**) qRT-PCR was used to measure *Tmprss2* mRNA in control and 2-month-old mice exposed to room air or hyperoxia and vehicle or mitoTEMPO between PND0-4 N=4-5 mice per group. Statistical significance is comparisons for all pairs using Tukey-Kramer HSD test with *P≤0.05; **P≤0.01.

## DISCUSSION

The COVID-19 outbreak was first detected in the Chinese city of Wuhan in 2019 and has since expanded rapidly to become one of the worst pandemics to ever challenge the modern world. While people of all ages are susceptible to infection, the severity of disease is worse in people who are elderly or who have pre-existing health conditions including COPD, diabetes, hypertension, and cancer ^28^. Those with multiple co-morbidities have a higher rate of mortality. People born preterm may also be at great risk for COVID-19 because they often suffer from multiple co-morbidities due, in part, to their lungs being exposed to oxygen too soon or to super-physiological concentrations used to maintain appropriate blood oxygen saturations. It is unclear whether co-morbidities increase disease by changing spatial and temporal expression of SARS-CoV-2 receptors or the immune response that leads to a lethal cytokine storm ^1^. In this study, we present evidence that expression of the SARS-CoV-2 co-receptors ACE2 and TMPRSS2 increase in the respiratory epithelium of mice as they age and this can be stimulated or accelerated by early exposure to hyperoxia. Expression of ACE2 in distal AT2 cells was of particular interest because infection of these cells with other viruses has been associated with higher mortality in humans ^15-17^. When infected such as by influenza A virus, AT2 cells may contribute to lung disease by producing inflammatory molecules that contribute to a lethal cytokine storm ^12^. They may also die and therefore reduce the number of surviving AT2 cells required to serve as stem cells for alveolar regeneration ^22,29,30^. Our findings support the idea that age and co-morbidities like preterm birth may increase the severity of COVID-19 by changing temporal and spatial patterns of SARS-CoV-2 receptors.

We found that ACE2 was primarily expressed by airway Club cells during early postnatal life. The intensity of ACE2 staining increased in the airways of mice with age and became detectable in the alveoli of young adult mice. Co-localization with proSP-C revealed that most, but not all alveolar cells expressing ACE2 were AT2 cells. Our findings are consistent with an earlier study showing that ACE2 is expressed in the adult mouse lung by Clara cells (now called Club cells), AT2 cells, and to some extent by endothelial cells around small and medium sized vessels ^31^. While that study showed how ACE2 levels rise during fetal development, our findings extend it by showing that ACE2 expression continues to increase as mice age. We also found that *Tmprss2* mRNA expression increases as mice age and this expression was similarly enhanced by neonatal hyperoxia. While AT2 cells have previously been shown to express TMPRSS2 ^11^, we were not able to detect it in the mouse lung using commercially available antibodies. However, we did find that the abundance of *Tmprss2* mRNA and protein abundance increased with age and neonatal hyperoxia, and was reduced by mitoTEMPO similar to that of *Ace2*. The higher expression of these genes as mice age is in agreement with recent review that discussed two unpublished studies deposited in *bioRxiv* showing how expression of *Ace2* and *Tmprss2* mRNA increases with age in human respiratory epithelium ^1^. Those findings in humans and ours in mice suggest the age-dependent increase in SARS2-CoV-2 receptors may be responsible for increasing the severity of COVID-19 lung disease in elderly people.

It is important to recognize the normal functions of ACE2 and TMPRSS2 because that may help explain why their expression steadily increases with age ^32^. ACE2 is perhaps best known for its role in controlling blood pressure in the renin-angiotensin system ^33^. ACE1 converts the 10-amino acid angiotensin I to an 8-amino acid vasoconstrictive peptide called angiotensin II. ACE2 accumulates in people with pulmonary hypertension and hydrolyzes Angiotensin II to Ang(1-7), which has vasodilation properties. Over-expressing ACE2 also protects against right ventricular hypertrophy ^34^. Hence, higher levels of ACE2 seen as the lung ages may reflect an adaptive response designed to protect against the development of cardiovascular disease. Interestingly, ACE2 levels decline in bleomycin-induced lung fibrosis and humans with interstitial pulmonary fibrosis while angiotensin II levels rise ^35,36^. Angiotensin II can promote fibrosis by stimulating AT2 cell apoptosis downstream of TGF-β signaling ^37^. ACE2 serves as an anti-fibrotic molecule by stimulating the hydrolysis of angiotensin II to Ang(1-7), which in turn signals through the Mas oncogene to block AT2 cell apoptosis by suppressing JNK activation ^38^. The slow and steady increase in ACE2 expression as the lung ages may also serve to preserve AT2 cells and thus reduce or prevent the development of idiopathic pulmonary fibrosis. In contrast to ACE2, the normal role of TMPRSS2 in the lung is poorly understood. TMPRSS2 is a serine protease that is localized to the apical surface of secretory cells such as Club and AT2 cells of the lung ^39^. Its expression is highly regulated by androgens in the prostate gland and may be similarly responsive to androgens in the lung, suggesting it may play a role in sex-dependent differences in the lung.

Our study also found that neonatal hyperoxia increased or accelerated expression of *Ace2* mRNA, ACE2 protein, and *Tmprss2* mRNA as mice age. Significant changes were seen with 60% or more FiO_2_ at 8 weeks (2 months) of age and persisted as mice age. How hyperoxia regulates expression of these proteins is conflicting and remains to be better understood. One study using human fetal IMR-90 fibroblasts found that hyperoxia does not change expression of ACE2 ^40^. However, ACE2 was depleted when cells returned to room air presumably because it was being proteolyzed and shed into the media. In contrast, another study found higher levels of ACE2 in newborn rats exposed to 95% oxygen for the first week of life and then recovered in 60% oxygen for the next two weeks ^41^. In our hands, changes in *Ace2* or *Tmprss2* mRNA were first detected in 8-week-old mice exposed to hyperoxia between PND0-4. We did not detect changes at the end of oxygen exposure (PND4). In fact, we recently deposited an RNA-seq analysis of AT2 cells isolated from PND4 mice exposed to room air versus hyperoxia that shows hyperoxia modestly inhibits *Ace2* and increases *Tmprss2* mRNA abundance (Gene Expression Omnibus of the National Center for Biotechnology Information under the accession number GSE140915). This suggests neonatal hyperoxia may not affect expression until after the mice are returned to room air. Because ACE2 and TMPRSS2 were only affected by doses of oxygen that cause long-term changes in lung function (i.e., 60% for 4 days but not 40% for 4 or 8 days), we speculate that they occur as an adaptive response to the alveolar simplification and cardiovascular disease as mice exposed to neonatal oxygen age. The elevated expression of ACE2 and perhaps TMPRSS2 may serve to prevent the loss of AT2 cells damaged by early oxygen and promote vasodilation as the pulmonary capillary bed undergoes rarefaction ^23,42^. But higher levels of these proteins may become a maladaptive response when they render the lung more susceptible to coronavirus infections.

While it remains to be determined how age or oxygen regulate expression of ACE2 and TMPRSS2, our studies with mitoTEMPO suggest their expression may be influenced by oxidative stress. Administering mitoTEMPO, a scavenger of mitochondrial superoxide during hyperoxia blunted the oxygen-dependent increase in these genes detected in 2-month-old mice. Because hyperoxia progressively increases mitochondrial oxidation, it has historically been used to model aging-related oxidative stress ^43^. This implies mitochondrial oxidation that accumulates as the lung ages steadily increases expression of ACE2 and TMPRSS2, which in turn may then attempt to defend against the pathological changes attributed to the aging process. Anti-oxidant therapies may therefore prove useful for suppressing expression of SARS-CoV-2 receptors and reducing the severity of COVID-19 related lung disease, especially in people with pre-existing co-morbidities.

Increased expression of ACE2 and TMPRSS2 may not be the only way these proteins enhance the severity of COVID-19-related lung disease. For example, TMPRSS2 facilitates viral activation and entry by cleaving hemagglutinin on influenza A virus and the spike protein on the SARS-CoV-2 virus ^11^. The spike protein accesses the cell when it binds the glucose regulated protein 78 (Grp78, BiP) found on the cell surface ^44^. Grp78 is a master regulator of the unfolded protein response (UPR) ^45^. It is normally localized to the endoplasmic reticulum (ER) where it inhibits the UPR by binding Activating Transcription Factor 6 (ATF6), Protein kinase RNA-like Endoplasmic Reticulum Kinase (PERK), and Inositol-requiring Enzyme 1 (IRE1). Grp78 is released from these proteins when oxidation and other stressful conditions cause an accumulation of unfolded proteins. It can then escape the ER and traffic to the cell surface where it becomes available to bind the coronavirus S protein and facilitate viral entry. This information should raise great concern for people with familial forms of IPF caused by mutations in *SFTPC* and other genes that activate the UPR in AT2 cells ^46^. Genetic studies in mice suggest mutant forms of SP-C that activate the UPR are not sufficient by themselves to cause fibrotic lung disease. However, they can predispose the lung to fibrotic disease following viral infections ^47^. Familial forms of IPF that activate the UPR in AT2 cells may therefore accelerate the age-dependent susceptibility of AT2 cells to SARS-CoV-2 infections.

In summary, we found that neonatal hyperoxia increases or accelerates the age-dependent expression of ACE2 and TMPRSS2 in the airway and alveolar epithelium of mice. Understanding how expression of these proteins changes with age and in response to early life insults such as neonatal hyperoxia may provide new opportunities for reducing the severity of COVID-19 and other types of lung disease.

## MATERIALS AND METHODS

### Mice

C57BL/6J mice were purchased from the Jackson Laboratories and maintained as an inbred colony. Mice were exposed to room air (21% oxygen) as control or hyperoxia (100% oxygen unless otherwise stated) between birth and postnatal day (PND) 4 and then returned to room air ^19^. Dams were cycled every 24 hours to ensure that hyperoxia did not compromise their health. Some mice exposed to room air or hyperoxia were injected intraperitoneally with 0.7μg/g mitoTEMPO (Enzo Life Sciences, Farmingdale, NY) or vehicle (phosphate-buffered saline) on PND0, PND1, and PND2. All mice used in this study were of mixed sex and housed in a pathogen-free environment according to a protocol (UCAR2007-121E) approved by the University Committee on Animal Resources at the University of Rochester.

### Immunohistochemistry

Lungs were inflation fixed overnight in 10% neutral buffered formalin, embedded in paraffin and 4 μm sections prepared ^23,48^. Sections were stained with antibodies against ACE2 (Invitrogen, PA5-47488, Waltham, MA), Scgb1a1 (Sigma, 07-063, St. Louis, MO) and proSP-C (Seven Hills Bioreagents, Cincinnati, OH). Immune complexes were detected with fluorescently labeled secondary antibody (Jackson Immune Research, West Grove, PA). Sections were then stained with 4’, 6-diamidino-2-phenylindole (DAPI) (Life Technologies, Carlsbad, CA) before viewing with Nikon E800 Fluorescence microscope (Microvideo Instruments, Avon, MA) and a SPOT-RT digital camera (Diagnostic Instruments, Sterling Heights, MI).

### Quantitative RT-PCR

Total RNA was isolated from the lung using Trizol reagent (ThermoFisher Scientific) and reverse transcribed to cDNA using the iScript cDNA synthesis kit (Bio-Rad Laboratories, Hercules, CA). The cDNA was then amplified with SYBR Green I dye on CFX96™ or CFX384™ Real-Time PCR detection system (Bio-Rad Laboratories, Hercules, CA). PCR products were amplified with sequence-specific primers for mouse *Ace2* (sense 5’-GGATACCTACCCTTCCTACATCAGC-3’ and antisense CTACCCCACATATCACCAAGCA-3’), *Tmprss2* (sense 5’-TACTTGGAGCGGACGAGGAA-3’, and antisense 5’-AGGAGGTCAGTATGGGGCTT-3’) or 18S rRNA (sense 5-CGGCTACCACATCCAAGGAA-3’, and antisense 5’-GCTGGAATTACCGCGGCT-3’) used to normalize equal loading of the template cDNAs. Amplifications were conducted with iTaq Universal SYBR Green Master Mix (Bio-Rad Laboratories, Hercules, CA). Fold changes in gene expression were calculated by the ΔΔCt method using the Ct values for the housekeeping 18S rRNA as a control for loading.

### Western blot analysis

The left lung lobe wsd homogenized in lysis buffer and insoluble material removed by centrifugation ^23^. Equal amounts of protein were separated on sodium dodecyl sulfate polyacrylamide gels and transferred to nylon membranes. The membranes were immunoblotted with primary antibodies to ACE2 (Invitrogen, PA5-47488, Waltham, MA), TMPRSS2 (Abcam, ab92323, Cambridge, MA) or β-ACTIN (Sigma, A2066). The blots were then incubated in appropriate secondary antibody (Southern Biotech, Birmingham, AL). Immune complexes were detected by chemiluminescence and visualized with a ChemiDoc Imaging System (Bio-Rad Laboratories, Hercules, CA).

### Statistical Analysis

Data were evaluated using JMP14 software (SAS Institute, Cary, NC) and graphed as means ± SEM. An unpaired t-test and 2-way ANOVA were used to determine overall significance, followed by Tukey-Kramer HSD tests.

## ACKNOWLEDGEMENTS

We thank Robert Gelein for maintaining the oxygen exposure facility and Daria Krenitsky for tissue processing and sectioning. This work was funded, in part, by National Institutes of Health Grants R01 HL091968 (M. A. O’Reilly), NIH Center Grant P30 ES001247, and a generous startup package from the Department of Pediatrics (A. M. Dylag). The University of Rochester’s Department of Pediatrics provided financial support through the Perinatal and Pediatric Origins of Disease Program.

## AUTHOR CONTRIBUTIONS

M.Y. designed and conducted experiments, analyzed the data, prepared figures, and helped write the manuscript; E.C. performed experiments and helped write the manuscript; J.H., performed experiments; A.D. aided in experimental design; M.O. designed the experimental research, analyzed the data, and wrote the manuscript. All authors reviewed and approved the final version of the manuscript.

